# Near chromosome-level genome assembly of the microsporidium *Hamiltosporidium tvaerminnensis*

**DOI:** 10.1101/2023.06.02.543461

**Authors:** Pascal Angst, Jean-François Pombert, Dieter Ebert, Peter D. Fields

**Author notes:** Corresponding author Pascal Angst, Department of Environmental Sciences, Zoology, University of Basel, Vesalgasse 1, 4051 Basel. Shared last author (Dieter Ebert, Peter D. Fields).

## Abstract

Microsporidia are intracellular parasitic fungi whose genomes rank among the smallest of all known eukaryotes. A number of outstanding questions remain concerning the evolution of their large-scale variation in genome architecture, responsible for genome size variation of more than an order of magnitude. This genome report presents a first near-chromosomal assembly of a large-genome microsporidium, *Hamiltosporidium tvaerminnensis*. Combined Oxford Nanopore, Pacific Biosciences, and Illumina sequencing led to a genome assembly of 17 contigs, 11 of which represent complete chromosomes. Our assembly is 21.64 Mb in length, has an N50 of 1.44 Mb, and consists of 39.56 % interspersed repeats. We introduce a novel approach in microsporidia, PacBio Iso-Seq, as part of a larger annotation pipeline for obtaining high-quality annotations of 3,573 protein-coding genes. Based on direct evidence from the full-length Iso-Seq transcripts, we present evidence for alternative polyadenylation and variation in splicing efficiency, which are potential regulation mechanisms for gene expression in microsporidia. The generated high-quality genome assembly is a necessary resource for comparative genomics that will help elucidate the evolution of genome architecture in response to intracellular parasitism.

**Significance:** Microsporidia are a model for genome evolution in response to intracellular parasitism, but we lack high-quality resources from species with large genomes. We present a near complete assembly of a large-genome microsporidium, *Hamiltosporidium tvaerminnensis*, and obtain high-quality gene annotations through full-length transcripts using Iso-Seq, a novel approach in microsporidia. Our study provides insights into gene regulation and paves the way for comparative genomic analyses aiming to understand the evolution of genome reduction and expansion in these intracellular parasites.

## Introduction

Microsporidia have become a valuable model in evolutionary biology for studying extreme parasitism (Murareanu et al., 2021; Wadi & Reinke, 2020). Compared to their fungi relatives, the genomes of microsporidia have been strongly shaped by their specialized life history of obligate intracellular parasitism (Corradi, 2015). In particular, the genomes of most microsporidia are very compact and reduced (Murareanu et al., 2021). However, genome sizes of microsporidia vary by more than an order of magnitude, making them a suitable model for studying genome evolution in eukaryotes and parasites (Wadi & Reinke, 2020). A powerful approach for this is comparative genomics (i.e., comparing the presence or absence of genetic sequences, their relative location, and their abundance among different species) for which high-quality genome assemblies are a prerequisite (Kille et al., 2022). Increasing interest in microsporidia has boosted available genome resources, but most published genomes are drafts (Williams et al., 2022). Following from recent successful efforts to obtain telomere-to-telomere assemblies in *Encephalitozoon*, a genus of microsporidia with small genomes (Mascarenhas dos Santos et al., 2023), we focus here on a large-genome species, *Hamiltosporidium tvaerminnensis*.

First noted by Green (1957) as *Octosporea bayeri*, the reclassified *H. tvaerminnensis* has become a model in the study of host–parasite interactions with its only host *Daphnia magna*, a planktonic microcrustacean (Altermatt et al., 2007; Lass & Ebert, 2005; Orlansky & Ben-Ami, 2019; Routtu & Ebert, 2015; Santos & Ebert, 2022; Vizoso & Ebert, 2004). Haag et al. (2011) provided the cytological and molecular description of *H. tvaerminnensis*, and Corradi et al. (2009) produced the first draft of its genome. Recently, genetic and genomic studies have started using *H. tvaerminnensis* to investigate the evolution of variation in microsporidian genome architecture (Angst et al., 2022, 2023; de Albuquerque et al., 2020; Haag et al., 2013, 2020). *Hamiltosporidium tvaerminnensis* has a remarkably large genome compared to other microsporidia, partly due to the proliferation of transposable genomic elements (de Albuquerque et al., 2020; Parisot et al., 2014). Draft genomes are available for the nearest known relative, *H. magnivora*, and more distantly related species, *Thelohania contejeani* (Cormier, Wattier, et al., 2021) and *Edhazardia aedis* (Desjardins et al., 2015).

Demands for comparative genomic approaches are increasing, calling for an improved genome of *H. tvaerminnensis*, a leading model organism among large-genome microsporidia. A high-quality genome will allow advancements in the study of genome architecture evolution in microsporidia; for example, through comparison with chromosomal assemblies of *Encephalitozoon*. We have integrated Pacific Biosciences (PacBio) Continuous Long Reads (CLR), Oxford Nanopore Technologies (ONT), and Illumina DNA sequencing for a near-chromosomal assembly, and we employed PacBio Iso-Seq for high-quality gene annotation. The first use of Iso-Seq for a microsporidium provides insights into microsporidian alternative polyadenylation (APA) and splicing efficiency as potential regulation mechanisms of gene expression in reduced genomes.

## Results and Discussion

### Genome Sequencing and Assembly

We generated a genome assembly of *H. tvaerminnensis* based on PacBio CLR and ONT sequencing reads (total/N50: 17.53 Gb/1.54 Kb and 2.75 Gb/32.68 Kb, respectively). In a subsequent step, we polished this assembly using Illumina paired-end (PE) sequencing reads (total: 66 Gb). The final assembly consisted of 21.64 Mb in 17 contigs with an N50 of 1.44 Mb (Table 1, Figure 1). We used BUSCO to assess the biological completeness of our assembly and found a BUSCO score of 94 %, a relatively high value for a large-genome microsporidium (Cormier, Chebbi, et al., 2021). The structural completeness of our assembly was assessed using scripts from (Mascarenhas dos Santos et al., 2023), which allow for the identification of telomeres. Telomeres are identifiable by terminal, short, tandem repeats– (TTAGGG)_*n*_ in the case of *H. tvaerminnensis*, as is the case in most fungi (Rahnama et al., 2020). Eleven contigs represented complete chromosomes flanked by telomeres at both ends (Figure 1, Table S1), four contigs had only one telomere, and two had none. Therefore, we estimated the genome contains 13 – 17 chromosomes. Large multicopy repeats arrayed in tandem and scattered throughout the genome prevented us from fully assembling the chromosomes with major repetitive loci abutting contig ends without telomeres (Figure 1). Overall, 39.56 % of the genome consisted of interspersed repeats (retroelements 7.07 %, DNA transposons 7.04 %, other 25.45 %). High numbers of repeats are common in large-genome microsporidia (de Albuquerque et al., 2020), explaining why previous attempts to assemble the genome of *H. tvaerminnensis* have been less successful in both completeness and contiguity (size = 18.34 Mb; number of contigs = 2,915; N50 = 9.58 Kb; BUSCO score = 87.6 %) (Haag et al., 2020) compared to our long-read sequencing-based assembly. The highest quality genomes within the microsporidia are from the genus *Encephalitozoon*, which have an order of magnitude smaller genomes than *H. tvaerminnensis* and telomeric repeats of (TTAGG)_*n*_ on both ends of all eleven chromosomes (Mascarenhas dos Santos et al., 2023).

**Table 1.**
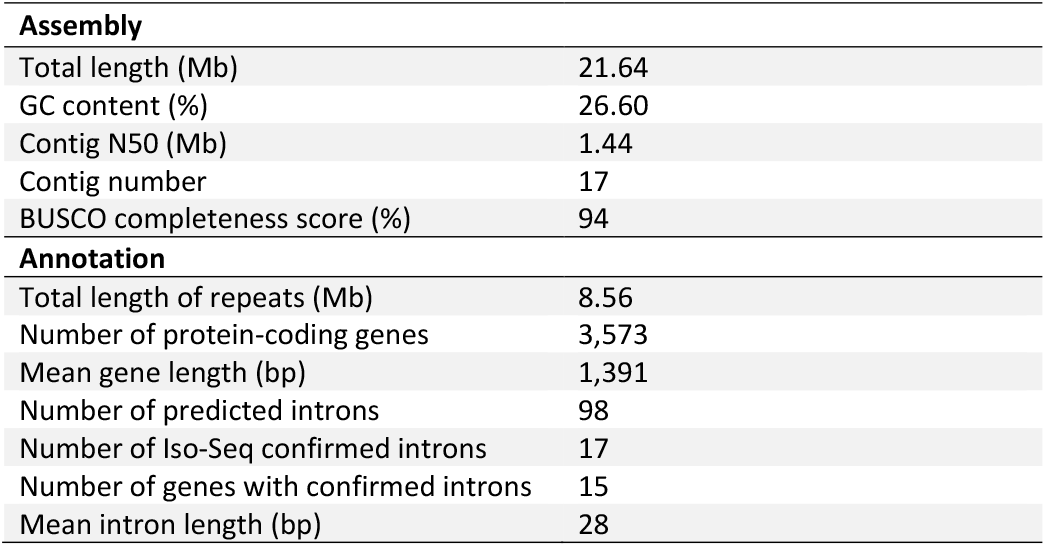
Statistics of the assembly and annotation of the *H. tvaerminnensis* genome.

**Figure 1.**
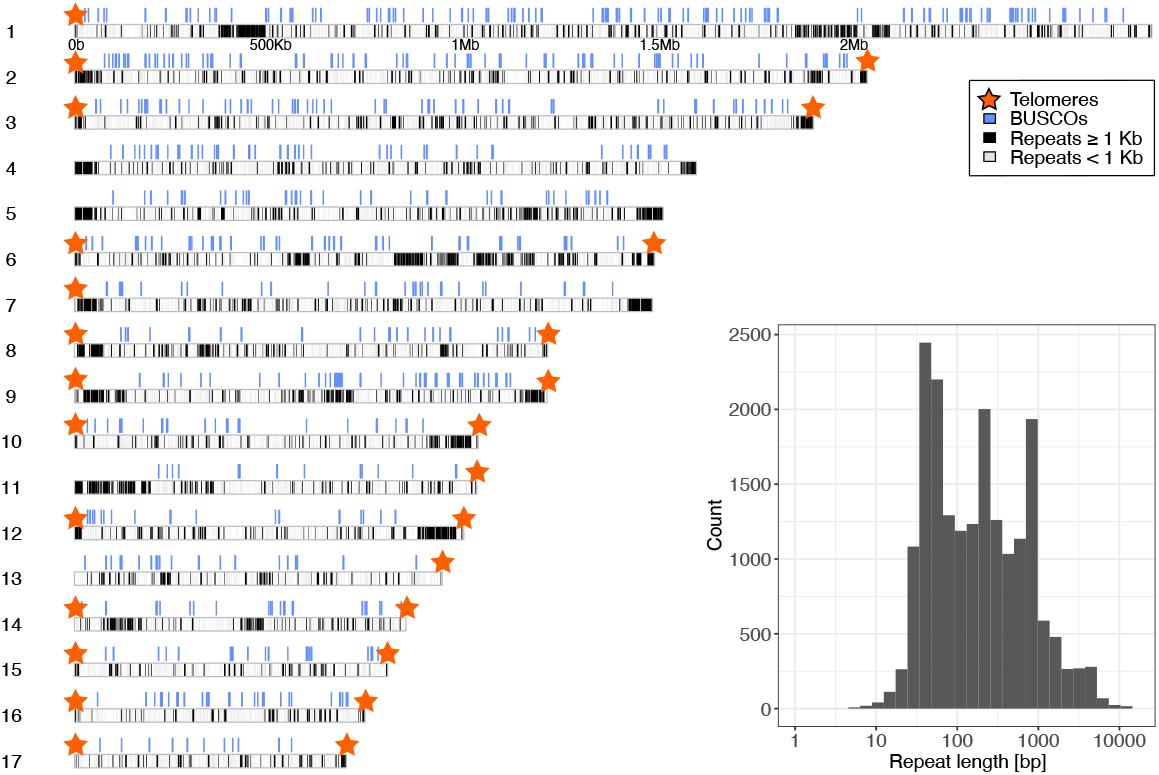
Karyoplot of *H. tvaerminnensis*. Eleven of the 17 contigs have two telomeres, i.e., represent full chromosomes. In total, 39.56 % of the genome are interspersed repeats and low complexity DNA sequences. The inset shows the repeat length distribution on a base-10 logarithmic scale.

### Iso-Seq for Genome Annotation

RNA-seq data from intracellular parasites are sparse, and even more sparse are full-length transcript sequences. Iso-Seq helped us to predict both gene structures and untranslated regions (UTRs) more accurately than was previously done for microsporidia. Using 8.5 Gb of Iso-Seq reads (so-called circular consensus sequence (CCS) of cDNA), we obtained annotations for 3,573 protein-coding genes with a mean length of 1,391 bp (Table 1). The number of genes in microsporidia ranges from about 2,000 to 4,000 and is positively correlated with genome size (Jespersen et al., 2022). Compared to previous work, we obtained fewer fragmented genes; the score of fragmented BUSCOs in our annotations was 1.8 % (previously 3.8 %; Haag et al., 2020). We also obtained many fewer introns (98) than previously predicted (890), but our number is still higher than estimates for other large-genome microsporidia annotated with the help of (short-read) RNA-seq (Desjardins et al., 2015). After observing the discrepancy between annotations with and without RNA-seq data, we suspected introns may not be efficiently spliced.

### Splicing Efficiency Analysis

Species with reduced genomes, sparse introns or no introns, and reduced spliceosomes have been suggested to exhibit low splicing efficiencies (Schärfen et al., 2022; Whelan et al., 2019). Microsporidia are such species. For example, in *E. cuniculi*, splicing efficiency for small introns is only around 20 % (Grisdale et al., 2013). We used direct evidence from full-length transcripts and found high variation in splicing efficiency in *H. tvaerminnensis* as has previously been reported for the reduced genomes of the red alga *Cyanidioschyzon merolae* (Schärfen et al., 2022) and for parasitic fungi (Sieber et al., 2018). We used 17 introns from 15 genes with good representation in the Iso-Seq data (Table S2). These introns had a mean length of 28 bp. The percentages of spliced transcripts ranged from 7.5 % to 92.3 % (Table S2) with a mean of 54.53 %. We did not find a statistical correlation between percent spliced and the length of introns. In conclusion, we found fewer introns in *H. tvaerminnensis* than previously suggested and intermediate splicing efficiencies, which is in line with theorized patterns for reduced-genome species. All but two introns identified in *H. tvaerminnensis* featured the same splice sites as have been found in (non-)parasitic fungi (i.e., 5’GU…AG3’) (Kupfer et al., 2004). The other two introns had splice sites 5’AA…AG3’ and 5’AU…GA3’.

### Alternative Polyadenylation

The mechanisms of gene regulation in microsporidia are unexplored. APA is a widespread and conserved regulatory mechanism by which the availability of multiple polyadenylation sites can lead to variation in 3’-UTR lengths of mRNA isoforms and thus 3’-UTR content, which strongly influences gene expression (Mitschka & Mayr, 2022). Here, Iso-Seq data allowed us to investigate the occurrence of mRNA isoforms encoding the same protein but differing in their 3’-UTRs resulting from APA. By observing multiple 3’-ends for many genes, we found direct evidence for APA in the full-length transcripts of *H. tvaerminnensis*. Upstream of the 3’-ends, a motif analysis of the 500 most expressed genes revealed enrichment of an UAAA tetramer. The UAAA tetramer was found in 95.8 % of the genes and 87.7 % of them were part of an A[A/U]UAAA hexamer on average 20 nucleotides upstream of the most common 3’-ends. A[A/U]UAAA is a known motif for cleavage of protein-coding mRNAs in metazoa (Neve et al., 2017) and is shown to be enriched upstream of 3’-ends in the bee-infecting microsporidium *Nosema ceranae* (Chen et al., 2021). These motifs were slightly less enriched upstream of the second and third most common 3’-ends (Table S3). No motif was strongly enriched that did not contain an UAAA tetramer.

## Materials and Methods

### Sample preparation and sequencing

The *H. tvaerminnensis* isolate used in this study was obtained from the *D. magna* genotype FI-OER-3-3, collected in July 2005 from a rockpool (59°47’18.9”N 23°10’26.9”E) on the Island Oeren in the Tvaerminne archipelago, Southwestern Finland. The same isolate has been Illumina sequenced for earlier genome drafts (Corradi et al., 2009; Haag et al., 2020) and a biogeographic study (Angst et al., 2022). Genomic Illumina reads from the latter were reused here (NCBI database, SRA accession: SRX13146514, Bioproject ID: PRJNA780787). We generated long-read sequencing data by applying a protocol described in Fields et al., (2015) for obtaining microbiota-free *D. magna* tissue, but retaining microsporidia, from which we extracted high-molecular-weight DNA using Genomic-tips (QIAGEN). A standard PacBio genomic DNA library and sequencing on a SMRTcell in CLR mode on a Sequel I have previously been done at the Quantitative Genomics Facility service platform at the Department of Biosystem Science and Engineering (D-BSSE, ETH) Basel, Switzerland (NCBI database, SRA accession: SRR23724335, Bioproject ID: PRJNA778105). Additionally, a ONT genomic DNA library was prepared with the SQK-LSK110 ligation kit and sequenced on a MinION device with a Spot-ON Flow Cell (R9.4.1).

To extract full-length transcripts, we used the RNeasy extraction kit (QIAGEN) supplying Proteinase K for protein digestion. The NEBNext Single Cell/Low Input cDNA Synthesis and Amplification Module kit (New England Biolabs) was used to generate cDNA from the extraction. An Iso-Seq cDNA library was generated with the Iso-Seq Express Oligo Kit and the SMRTbell Express Template Prep Kit 2.0 (PacBio). The Iso-Seq library was sequenced using a single SMRTcell on a Sequel I at the D-BSSE (Basel, Switzerland) and processed with the *isoseq3* pipeline (https://github.com/PacificBiosciences/IsoSeq), resulting in approximately 8.5 Gb of CCS data.

### Genome assembly

An assembly based on the PacBio CLR sequencing data, generated with canu v.2.1.1 (Koren et al., 2017), was extended *in-silico* using a seed-based approach with Consed v.29 (Gordon et al., 1998) as described in Pombert et al. (2013). Independently, the Nanopore sequencing reads were basecalled with bonito v.0.5.3 (https://github.com/nanoporetech/bonito) using the dna_r9.4.1_e8.1_sup@v3.3 model and then assembled with nextDenovo v.2.5.0 (https://github.com/Nextomics/NextDenovo) using an expected genome size of 20 Mb. The Nanopore nextDenovo assembly was polished with the Illumina reads and the PacBio CLR reads using nextPolish v.1.4.1 (Hu et al., 2020), Medaka v.1.7.2 (https://github.com/nanoporetech/medaka), and two runs of Pilon v.1.24 (Walker et al., 2014) and then merged with the extended PacBio CLR assembly using quickmerge v.0.3 (Solares et al., 2018) and manual curation. The absence of contaminants in this final assembly was ascertained by BLAST homology searches against the NCBI database and its completeness was assessed with check_for_telomeres.pl v.0.3 (Mascarenhas dos Santos et al., 2023). Repeats in the assembly were detected with RepeatModeler v.2.0.2 including the LTR pipeline (Flynn et al., 2020) and ReapeatMasker v.4.1.2 (Smit et al., 2013) and plotted with karyoploteR v.1.24.0 (Gel & Serra, 2017) in R v.4.2.2 (R Core Team, 2022).

### Genome annotation

The *H. tvaerminnensis* genome was annotated by feeding the Iso-Seq transcriptome data as extrinsic evidence to the fungal annotation pipeline funannotate v.1.8.13 (Palmer & Stajich, 2022). Briefly, as per guideline for gene prediction using long RNA reads (https://github.com/Gaius-Augustus/BRAKER/blob/master/docs/long_reads/long_read_protocol.md), we mapped the whole-transcript reads from the CCS FASTA produced by *isoseq3* to the repeat-masked genome using Minimap2 v.2.22 (Li, 2018) with the flags ‘-ax splice –secondary=no -C5’. Approximately 25.5 % of the reads aligned to the parasite genome. We then collapsed the total read set into distinct transcripts (while retaining isoforms) using the *collapse_isoforms_by_sam*.*py* script provided by the *cDNA_Cupcake* (https://github.com/Magdoll/cDNA_Cupcake) pipeline and ran GeneMarkS-T v.5.1 (Tang et al., 2015) to predict protein-coding regions in the transcripts. The resultant FASTA and GFF files were used for gene prediction and annotation with the wrapper tool funannotate v.1.8.13 (Palmer & Stajich, 2022), which relies on TANTAN v.26 (Frith, 2011) for repeat-masking. Funannotate operated AUGUSTUS v.3.4.0 (Stanke et al., 2006, 2008), GeneMark-ES v.4.62 (Brůna et al., 2020), GlimmerHMM v.3.0.4 (Majoros et al., 2004), SNAP v.2013_11_29 (Korf, 2004), and EVidenceModeler v.1.1.1 (Haas et al., 2008) for ab-initio gene prediction and consensus gene structure generation. Functional annotation in funannoate relies on the combined evidence from InterProScan v.5.55_88.0 (Jones et al., 2014), eggNOG-mapper v.2.1.9 (Cantalapiedra et al., 2021; Huerta-Cepas et al., 2019), Phobius v.1.01 (Käll et al., 2004), SignalP v.5.0b (Almagro Armenteros et al., 2019), and antiSMASH v.6.0 (Blin et al., 2021) as well as on comparisons to Pfam (Mistry et al., 2021), UniProt (The UniProt Consortium, 2022), MEROPS (Rawlings et al., 2014), dbCAN (Yin et al., 2012) and BUSCO v.5.4.3 (Manni et al., 2021) databases. Also, the completeness was assessed using BUSCO with its microsporidia_odb10 database (Creation date: 2020-08-05). For proteins without functional annotation from this pipeline, we lifted-over functional annotations from the species reference proteome (UniProt database, Proteome ID: UP000292282) using BLAST v.2.3.0+ (Camacho et al., 2009), AGAT v.1.0.0 (Dainat, 2022), and GFF3sort v.1.0.0 (Zhu et al., 2017).

### Full-length transcript analyses

For the analysis of splicing efficiency and APA, we followed the methodology described in Schärfen et al. (2022). Briefly, starting from the BAM file, which was provided by the sequencing facility, we used lima v.2.6.0 (https://github.com/PacificBiosciences/barcoding) in “-isoseq” mode to remove IsoSeq template-switching oligo sequences and to orient subreads in the correct 5’ to 3’ direction. After aligning subreads to our genome using minimap2 in “splice:hq” mode, we used SAMtools v.1.16.1 (Li et al., 2009) and BEDtools v.2.30.0 (Quinlan, 2014) to identify intron-spanning reads. The percentage of spliced introns was calculated by dividing the number of spliced reads by the total number of reads for each intron times 100. For this analysis, introns were manually selected based on Iso-Seq coverage level. A potential correlation between the percent of spliced reads and intron length was tested in R.

For the analysis of APA, briefly, we used SAMtools, BEDtools, and pysam v.0.20.0 (https://github.com/pysam-developers/pysam) to extract each read’s 3’-end coordinate following Schärfen et al. (2022). Then, we drew the distribution of 3’-end coordinates as a histogram separately for each gene and applied the find_peaks algorithm from scipy.signal v.1.7.3 (Virtanen et al., 2020) with height equal to four and distance equal to 18. Hence, we found the number of peaks (=3’-end coordinates) per gene. We then separately extracted sequences of the 500 most expressed genes up to 40 nucleotides upstream of the highest peak, the second highest peak, and the third highest peak using seqtk v.1.3-r106 (https://github.com/lh3/seqtk). For each set of sequences, we did a motif analysis using XSTREME v5.5.0 from the MEME Suite (Grant & Bailey, 2021) with a minimum motif length of four and determined the distance of the obtained motif from the peak.

## Data Availability Statement

Raw data is deposited at the NCBI SRA database and the assembled genome as well as predicted set of protein sequences are available at NCBI GenBank (FIOER33 v3; BioProject ID PRJNA778105) and at https://doi.org.10.6084/m9.figshare.22794887 and will be publicly available upon acceptance.

## Author Contributions

All authors designed the study. DE collected the sample. PA and PDF performed the molecular work. PA, JFP, and PDF analyzed the data. PA wrote the manuscript. All authors reviewed the manuscript.

## Acknowledgements

We thank Jürgen Hottinger, Urs Stiefel, Michelle Krebs, and Alix Thivolle for help in the laboratory. We thank members of the Ebert group for providing feedback on the study and the manuscript.

