## Supplemental Tables 1-3 for "Near chromosome-level genome assembly of the microsporidium *Hamiltosporidium tvaerminnensis*"

^*^ Corresponding author

**Table S1. Output from check_for_telomeres.pl.** For each scaffold, terminal, short, tandem repeats are reported based on which the presence of telomeres are determined, and chromosomal completeness is inferred.

| **Contig** | **Length** | **# of pattern (start)** | **# of rev pattern (start)** | **# of pattern (end)** | **# of rev pattern (end)** | **Status** |
| --- | --- | --- | --- | --- | --- | --- |
| scaffold_1 | 2,681,470 | 0 | 165 | 2 | 1 | partial chromosome with 1 telomere end |
| scaffold_2 | 1,973,229 | 0 | 139 | 115 | 1 | complete chromosome |
| scaffold_3 | 1,839,609 | 2 | 23 | 166 | 0 | complete chromosome |
| scaffold_4 | 1,547,981 | 1 | 0 | 1 | 1 | contig with no chromosome end attached |
| scaffold_5 | 1,464,916 | 0 | 2 | 2 | 0 | contig with no chromosome end attached |
| scaffold_6 | 1,444,059 | 0 | 145 | 127 | 0 | complete chromosome |
| scaffold_7 | 1,438,879 | 0 | 166 | 0 | 0 | partial chromosome with 1 telomere end |
| scaffold_8 | 1,177,745 | 1 | 81 | 57 | 0 | complete chromosome |
| scaffold_9 | 1,176,229 | 0 | 59 | 43 | 0 | complete chromosome |
| scaffold_10 | 1,005,214 | 1 | 95 | 145 | 0 | complete chromosome |
| scaffold_11 | 1,001,064 | 0 | 0 | 78 | 0 | partial chromosome with 1 telomere end |
| scaffold_12 | 969,012 | 2 | 79 | 111 | 0 | complete chromosome |
| scaffold_13 | 915,172 | 0 | 2 | 124 | 1 | partial chromosome with 1 telomere end |
| scaffold_14 | 824,917 | 0 | 117 | 56 | 0 | complete chromosome |
| scaffold_15 | 779,687 | 0 | 109 | 64 | 0 | complete chromosome |
| scaffold_16 | 723,150 | 0 | 68 | 115 | 1 | complete chromosome |
| scaffold_17 | 676,572 | 0 | 166 | 128 | 0 | complete chromosome |

**Table S2. Introns used in splicing efficiency analysis.**

| **Contig** | **Start Position** | **End Position** | **Gene ID** | **Strandedness** | **Fraction Spliced** |
| --- | --- | --- | --- | --- | --- |
| scaffold_4 | 140955 | 140975 | ID=FUN_3561;Parent=FUN_3561-T2 | - | 0.703125 |
| scaffold_4 | 141274 | 141296 | ID=FUN_3561;Parent=FUN_3561-T2 | - | 0.539683 |
| scaffold_11 | 223210 | 223247 | ID=FUN_002612;Parent=FUN_002612-T2 | - | 0.075085 |
| scaffold_4 | 299131 | 299152 | ID=FUN_3563;Parent=FUN_3563-T1 | - | 0.333333 |
| scaffold_12 | 302732 | 302759 | ID=FUN_002799;Parent=FUN_002799-T1 | + | 0.923077 |
| scaffold_11 | 410886 | 410946 | ID=FUN_002649;Parent=FUN_002649-T1 | - | 0.525 |
| scaffold_16 | 481659 | 481726 | ID=FUN_003286;Parent=FUN_003286-T1 | - | 0.653846 |
| scaffold_15 | 499747 | 499778 | ID=FUN_3480;Parent=FUN_3480-T2 | - | 0.384615 |
| scaffold_10 | 544104 | 544128 | ID=FUN_002544;Parent=FUN_002544-T2 | + | 0.638889 |
| scaffold_3 | 569729 | 569749 | ID=FUN_000955;Parent=FUN_000955-T1 | - | 0.307692 |
| scaffold_15 | 575854 | 575875 | ID=FUN_3482;Parent=FUN_3482-T2 | + | 0.5 |
| scaffold_10 | 629004 | 629028 | ID=FUN_002560;Parent=FUN_002560-T1 | - | 0.724138 |
| scaffold_5 | 731097 | 731121 | ID=FUN_001542;Parent=FUN_001542-T2 | + | 0.615385 |
| scaffold_5 | 731175 | 731194 | ID=FUN_001542;Parent=FUN_001542-T2 | + | 0.615385 |
| scaffold_6 | 780269 | 780291 | ID=FUN_001777;Parent=FUN_001777-T1 | + | 0.428571 |
| scaffold_6 | 780708 | 780729 | ID=FUN_001777;Parent=FUN_001777-T1 | + | 0.785714 |
| scaffold_2 | 1330684 | 1330706 | ID=FUN_000723;Parent=FUN_000723-T1 | + | 0.516129 |

**Table S3. Sequence motifs enriched in 40 nucleotides (nt) upstream of the most common, second most common, and third most common 3’-ends (peaks) and their frequency in the 500 most expressed genes.**

| **Motif** | **Frequency upstream of highest peak** | **Frequency upstream of second highest peak** | **Frequency upstream of third highest peak** | **Median distance between motif and highest peak (nt)** |
| --- | --- | --- | --- | --- |
| A.UAAA | 434 | 232 | 220 | 20 |
| AAUAAA | 260 | 130 | 131 | 20 |
| AUUAAA | 160 | 92 | 78 | 20 |
| UAAA | 479 | 368 | 360 | 18 |
